# Deep learning-enabled speckle reduction for cleared-sample coherent scattering tomography

**DOI:** 10.64898/2026.01.27.702188

**Authors:** Chong Chen, Huiru Wang, Peilin Gu, Xi Chen, Jian Ren

## Abstract

Clearing Assisted Scattering Tomography (CAST) extends coherent scattering tomography to whole-brain imaging, enabling visualization of fine-scale brain-wide connectivity. As a coherent optical tomography modality, CAST is inherently affected by speckle noise, which degrades image quality and limits quantitative analysis. However, existing speckle reduction methods developed for optical coherence tomography (OCT) are not directly transferable to CAST images due to differences in sample and noise statistics. Here, we present a learning-based cleared-sample speckle reduction network, termed CLEAR Net, specifically designed for CAST imaging, which effectively suppresses speckle noise in whole-brain white matter images while preserving fine structural details. We quantitatively benchmarked CLEAR Net against representative speckle reduction algorithms on CAST datasets and further evaluated its generalizability using publicly available ophthalmic datasets.

## 1. Introduction

Optical Coherence Tomography (OCT) has become an indispensable imaging modality in ophthalmology^1^, providing micrometer-scale structural visualization of retinal layers with exceptional sensitivity and a wide dynamic range. However, its application to large organs is fundamentally constrained by the limited optical penetration depth of biological tissue. It is generally recognized that when the imaging depth exceeds approximately 300 µm, multiply scattered photons begin to dominate the detected OCT signal, leading to severe contrast degradation and loss of structural fidelity^2^.

Tissue-clearing techniques have recently emerged as an approach to address the limited optical penetration depth in biological tissue. By reducing refractive-index mismatches and removing lipid components, tissue clearing substantially suppresses multiple scattering and enhances light penetration in thick specimens^3^. In our previous work, we developed a clearing assisted scatttering tomography (CAST) method, which combines tissue clearing with coherent scattering imaging to enable label-free, high-throughput volumetric imaging of intact mouse brains at micrometer resolution, markedly enhancing the visualization of fine structural features, including neural fibers and vasculature^4^. Nevertheless, despite the substantial reduction of multiple scattering, CAST images remain affected by residual speckle noise, particulary in small-scale structures. This speckle noise limits image uniformity and prevents consistently high-resolution reconstruction across the entire brain volume.

Researchers have proposed various methods to suppress speckle noise, which can be broadly categorized into hardware-based and software-based methods. Hardware-based methods reduce speckle noise by modifying the acquisition system to generate uncorrelated speckle patterns, such as angular compounding^5^, spatial compounding^6^, and frequency compounding^7^. Although such methods can reduce speckle, they require high system complexity, and the number of uncorrelated speckle patterns that can be generated is limited, making it impossible to completely eliminate noise. Moreover, since the acquisition methods of the compounded images differ, the reconstructed image often suffers from resolution loss. In addition, hardward based methods relies on the repeated acquisition of a large number of B-scans, significantly reducing imaging speed and temporal resolution^8^.

Software-based despeckle methods use algorithms to post-process acquired images, demanding much less data compared to hardware-based methods. Existing speckle reduction methods can be broadly categorized into temporal, spatial, and transform-domain approaches^9–11^.BM3D leverages block matching and three-dimensional transforms to preserve high-frequency features, yet may introduce artifacts and result in incomplete speckle reduction ^12^. Non-local filtering methods exploit image self-similarity beyond local neighborhoods. While Non-Local Means (NLM) and its probabilistic variant (PNLM) aggregate information from similar patches, PNLM relies on an additive Gaussian noise assumption that inadequately models speckle in coherent imaging^13^. More recently, TNode incorporates more realistic speckle statistics and three-dimensional voxel similarity search, leveraging inter-layer information to achieve state-of-the-art non-local despeckling performance while preserving tissue structures^14^. Despite their effectiveness, non-local filtering methods are computationally expensive due to exhaustive voxel-wise similarity searches, making them impractical for large-scale volumetric brain imaging, and their limited parameter flexibility often leads to oversmoothing of fine structures.

Owing to their large parameter space and strong representational capacity, deep learning–based methods have emerged as a dominant approach for speckle reduction, offering both high generalizability across diverse structural patterns and efficient inference for large-scale imaging data. Within this paradigm, both supervised and unsupervised learning strategies have been actively explored. Supervised methods leverage paired training data to learn direct mappings from noisy to clean images, while unsupervised and self-supervised approaches relax the requirement for paired datasets by exploiting statistical correlations or domain consistency^15–19^. Together, these approaches form a broad and rapidly evolving toolbox for speckle reduction in coherent imaging. However, existing deep learning-based speckle reduction methods have been developed almost exclusively using retinal OCT datasets, causing networks to implicitly learn structural priors and noise statistics specific to retinal tissue.To date, no studies have applied similar speckle reduction approaches to CAST brain imagng. This lack of transferability is expected, as retinal and brain tissues differ substantially in structural complexity, information content, and image entropy. Moreover, unlike conventional OCT images, which are predominantly affected by multiple scattering, CAST images are acquired from optically cleared samples. As a result, even identical anatomical structures exhibit markedly different speckle characteristics between OCT and CAST.

Based on the above challenges, our study aims to develop an efficient speckle reduction pipeline capable of preserving tissue details and rich structural information for whole-brain CAST images. In this work, We develop a supervised learning strategy to leverage physically meaningful paired CAST data, enabling more accurate modeling of speckle statistics and improved structural fidelity compared to unsupervised approache. We first constructed an improved algorithm based on TNode to generate high-fidelity silver ground truth. Building on this, we propose a GAN-based deep denoising framework that employs an attention-augmented backbone and a hybrid loss function to effectively guide image reconstruction. Experimental results demonstrate that the CLEAR Net achieves high fidelity in both visual appearance and quantitative metrics, effectively suppressing speckle while preserving fine brain structures, thereby providing a feasible solution for high-resolution whole-brain CAST images. CLEAR Net achieves speckle reduction performance comparable to hardware-based approaches and consistently outperforms TNode. In addition, CLEAR Net demonstrates measurable generalizability on ophthalmic OCT datasets.

## 2. Methods

### 2.1 Raw CAST Data Acquisition and Preprocessing

In the experiments, mouse brain tissues were cleared using a protocol based on the SWITCH method^20^, which was further optimized in our study^4^. We adjusted the clearing time and the refractive index matching conditions to improve tissue transparency, enhancing optical penetration and providing the best contrast in white matter regions. Mice were first perfused with PBS and then fixed with a solution of 4% paraformaldehyde (PFA) and 1% glutaraldehyde (GA). Following harvest, brains were incubated in the same fixation solution at 4 °C with gentle shaking for 3 days. Samples were then washed twice in PBST at room temperature, followed by overnight incubation in an inactivation solution (4% (w/v) acetamide and 4% (w/v) glycine) at 37 °C. Tissue clearing was performed in a solution of 200 mM sodium dodecyl sulfate (SDS) and 20 mM sodium sulfite, with incubation time adjusted according to sample size to achieve the desired optical transparency. After tissue clearing, the brains were immersed in exPROTOS refractive index-matching solution and incubated for 2 days.

Imaging was performed using our customized CAST system^4^. The homemade wavelength swept laser utilized a semiconductor optical amplifier (Covega Corp., BOA-4379) as the gain medium and a polygon scanner (Lincoln Laser Co., SA34/DT-72-250-025-AA/#01B) as the tunable filter to rapidly tune the wavelength at a rate of 54 kHz. This source had a center wavelength of 1300 nm and a sweeping range of 110 nm. The axial and lateral resolutions in air were approximately 9 µm, with an imaging depth of up to 11 mm. The reconstructed and stitched CAST volumetric images of whole brain had a resolution of 1280 × 3620 × 2048 (A-scan × B-scan × C-scan), of which 70% was used for training, 20% for validation, and 10% for testing. The raw data were interpolated and cropped to 2500 × 7240 × 4096. And then, the volume was divided into 512 × 512 patches along the *en-face* direction for training and validation. From the filtered input-GT pairs, 30,000 patches paired with the silver ground truth were randomly selected for training.

### 2.2 Ground Truth Images Construction

An improved TNode algorithm was employed to rapidly generate a large set of speckle-suppressed ground-truth images. The basic principle of TNode is that speckle noise in coherent imaging follows a Gamma distribution. The expectation of the exponential distribution corresponds to the true value, which can be reconstructed by averaging multiple independent samples. In the absence of repeated independent acquisitions, structurally similar but speckle-variant volume blocks within the tissue can be utilized. By computing similarities and applying weighted averaging, the true value can be estimated, thereby suppressing nearly independent speckle while preserving structure-related tissue information. This enables high-fidelity reconstruction of coherent imaging voxels.

In the TNode algorithm, similarity is calculated based on local window matching. Specifically, a similarity window is defined, and candidate blocks are searched within a designated search window. For each pair of windows, Bayesian likelihood is computed by pixel-wise intensity comparison, and the maximum likelihood estimate is used to approximate the true value^21^. This yields a generalized likelihood ratio, which is further defined as the metric for similarity evaluation *S*_*i*_*(x, y*_*i*_ ).

At the voxel level,

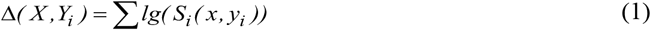

where *S*_*i*_(*x, y*_*i*_) denotes the similarity between two pixels and Δ(*X*,*Y*_*i*_) denotes the window-level similarity derived from pixel correlations.

Similarity weights are computed as

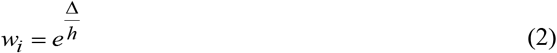

where the parameter *h* controls sensitivity to noise level.

The final reconstruction formula is given by:

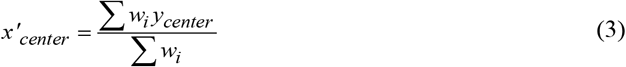

where 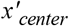 and *y*_*center*_ refer to the central pixel of the volume block in the reconstructed and Y windows respectively.

TNode relies on a similarity formulation that approximates the true signal only under relatively low-noise conditions. In retinal OCT imaging, where tissues are comparatively homogeneous and SNR is typically high, reconstruction can be effectively regulated by an SNR-dependent parameter *h* enabling robust speckle reduction while preserving structural information. However, under low-SNR conditions, the similarity assumption becomes unreliable, limiting the performance of TNode.In contrast, CAST brain images typically suffer from lower SNR and contain more intricate and fine tissue structures that are easily obscured or fragmented by noise. In such cases, relying solely on parameter *h* often leads to over-smoothing, causing the loss of fine structural details.

To address this issue, we introduced a similarity threshold into the TNode framework: only the top 10% most similar candidate blocks within the search window were retained for reconstruction. This more conservative strategy effectively reduces over-smoothing risk. Moreover, in CAST brain imaging, fine structures (e.g., white matter fibers) are close in size to speckle (approximately 4 pixels vs. 3 pixels, respectively). During parameter tuning of the modified TNode, we focused primarily on weight factors related to *h*, the radii of the similarity and search windows, and reduction thresholds for additive noise. The final optimal configuration was:

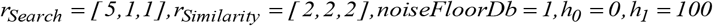

Although this setup produced superior reconstruction results, processing a full mouse brain volume still required about 6-7 days, motivating the use of deep learning to further enable high-throughput processing.

### 2.3 Network Architecture

To achieve efficient high throughput speckle-suppressed reconstruction, we designed a GAN-based framework. The GAN framework relies on adversarial training between a generator and a discriminator: the generator is optimized to produce outputs indistinguishable from the ground truth, while the discriminator is optimized to distinguish generated samples from real ones. Through this adversarial process, both networks improve simultaneously, allowing the generator to capture more realistic and detailed structural features.

The generator is a U-Net with four encoder-decoder layers and a bottleneck layer for further feature processing. Skip connections between encoder and decoder preserve shallow information. An attention mechanism was introduced into convolutional modules to selectively retain edge and texture features while suppressing noisy regions. The discriminator is a four-layer convolutional PatchGAN network. PatchGAN constrains realism within local receptive fields, guiding the generator to preserve details and avoid over-smoothing^22,23^. The generator takes a 512 × 512 speckle-contaminated CAST patch as input and outputs a denoised patch. The discriminator takes either an (input, generated) pair or an (input, GT) pair and outputs a prediction map labeling corresponding regions as real or fake.

### 2.4 Training Strategy and Loss Functions

To ensure that the learning rates of the generator and discriminator remain balanced, preventing one from learning too fast and failing to provide sufficient information for adversarial training, we adopted an alternating training strategy: two epochs of generator-only training followed by one epoch of joint generator-discriminator training.

During generator-only training, the loss function is:

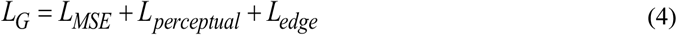

where *L*_*perceptual*_ is computed as the feature difference between GT and generated data using the VGG conv3-3 layer, ensuring high-dimensional texture consistency. *L*_*edge*_ preserves edge structures and prevents over-smoothing^24^. Perceptual and edge losses both help generator to maintain structural fidelity. SSIM loss was not used during training to avoid premature optimization toward evaluation metrics. In this stage, only generator parameters are updated, while the discriminator remains unchanged.

During joint generator-discriminator training, the generator loss is:

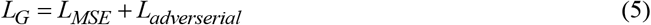

while the discriminator loss is:

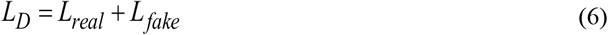

Both generator and discriminator parameters are updated simultaneously at this stage.

Training was conducted on a NVIDIA A100 GPU. Adam was used as the optimizer for both networks, with learning rates of 0.01 for the generator and 0.001 for the discriminator, and momentum parameters set to (0.5, 0.999).

### 2.5 Experiments and Evaluation Metrics

The method’s performance was evaluated using PSNR (pixel intensity difference), SSIM (structural similarity), EPI (edge preservation index) and CNR (contrast discrimination against noise). To be specific, evaluation metrics are defined by the following formulas:

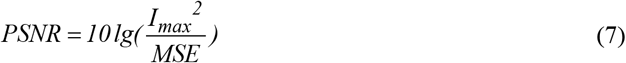

where *I*_*max*_ is the maximum value in the estimated image, and 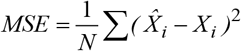 is the mean square error between the ground truth voxels and the estimated image voxels.

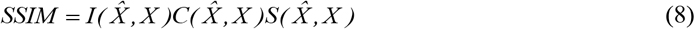

where 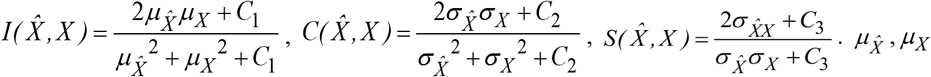 are the mean intensities of grounf truth image and estimated image. 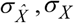 are their standard deviations and 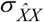 is their covariance. *C*_*1*_, *C*_*2*_ and *C*_*3*_ are small positive values to stabilize the computation.

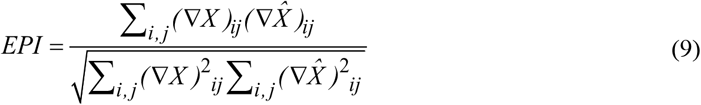

where ∇*X* and 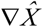 correspond to the per-pixel magnitude of the spatial gradients of the ground-truth and generated images respectively, and ε is a small constant to avoid numerical instability.

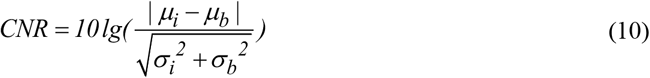

where *μ*_*i*_ and *μ*_*b*_ are the mean of region of interest (ROI) and background region respectively. σ _*i*_ andσ_*b*_ are the standard deviations of ROI and background region respectively.

We experimented with different network and training configurations, including the number of convolutional layers in the discriminator, real-label smoothing strategies, and learning rate settings, to determine the optimal architecture and loss function combination.

Comparative experiments included classical methods (PNLM, BM3D) as well as learning-based methods (Sub2Full^25^). Results generated by these methods were compared against clean reference images generated from speckle modulating or TNode, and quantitative evaluations were performed using PSNR, SSIM, EPI and CNR.

## 3. Results and Discussion

In this study, we used CAST-acquired mouse brain volumes and randomly extracted about 30,000 patches of size 512×512 for training from scratch, with a batch size of 8 and a total of 100 iterations. Experimental results demonstrate that the proposed GAN framework can effectively generate speckle-reduced volumes close to the Ground Truth, achieving significant improvements in quantitative evaluation metrics, while also greatly improving processing speed compared to the traditional algorithm used for Ground Truth generation.

### 3.1 CLEAR Net speckle reduction results of CAST brain images

Figure 2 shows the whole-brain CAST images after speckle reduction using CLEAR Net. A three-dimensional rendering of entire brain, togethre with two representative regions of interest (ROIs) from the cerebellum and cerebrum exhibiting distinct structural organization, are present in Fig.2b-c. As indicated by the white arrows and magnified views, the raw CAST contain substantial speckle nosie that obscures fine structural details. In contrast, the CLEAR Net processed images exhibit effective speckle reduction while preserving fine anatomical features in both gray and white matter regions. In the cerebral ROIs, CLEAR Net markedly improves texture continuity in white-matter-rich regions, revealing delicate fibrous patterns and enhancing overall structural clarity (Fig.2d-e). Three sub-ROIs (i, ii, and iii) are presented for a closer examination of specific structures. In the magnified pictures, raw images exhibit apparent speckle noise. After processing, the CLEAR Net processed images reveals enhanced contrast and more distinct delineation of fine structures. Overall, the results confirm that our pipeline not only generalizes well across different brain regions but also produces consistent, artifact-free reconstructions. The enhanced contrast and improved delineation of fine and layered structures demonstrates our pipeline’s potential for high-precision 3D image restoration and analysis.

**Figure 1.**
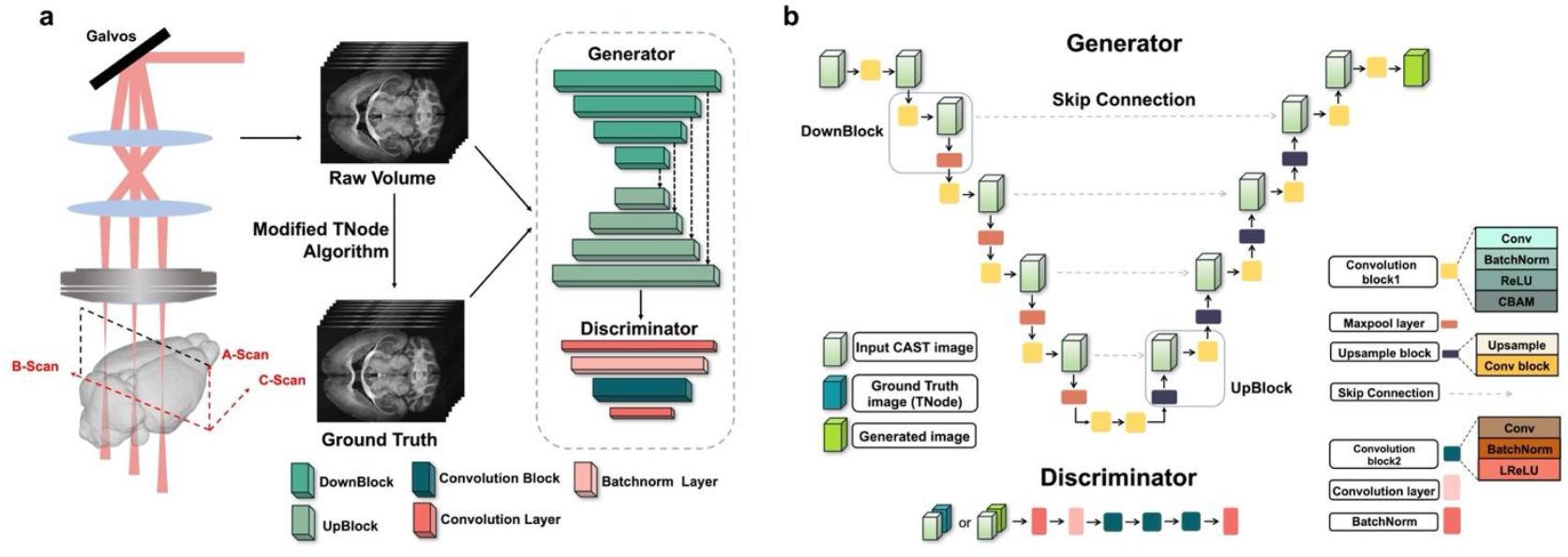
Overall pipeline and architecture of the proposed GAN-based CAST imaging speckle reduction framework, CLEAR Net. (a) Schematic of the scattering imaging and data preparation pipeline. Raw volumes are acquired using the CAST imaging system. The modified TNode algorithm is then applied to generate the Ground Truth for supervised training. (b) Architecture of the proposed GAN framework. The generator adopts a U-Net backbone with CBAM-enhanced convolutional blocks, integrating DownBlocks, UpBlocks, and skip connections for structure preservation. The discriminator follows the PatchGAN design to distinguish generated images from TNode-derived ground truths.

**Figure 2.**
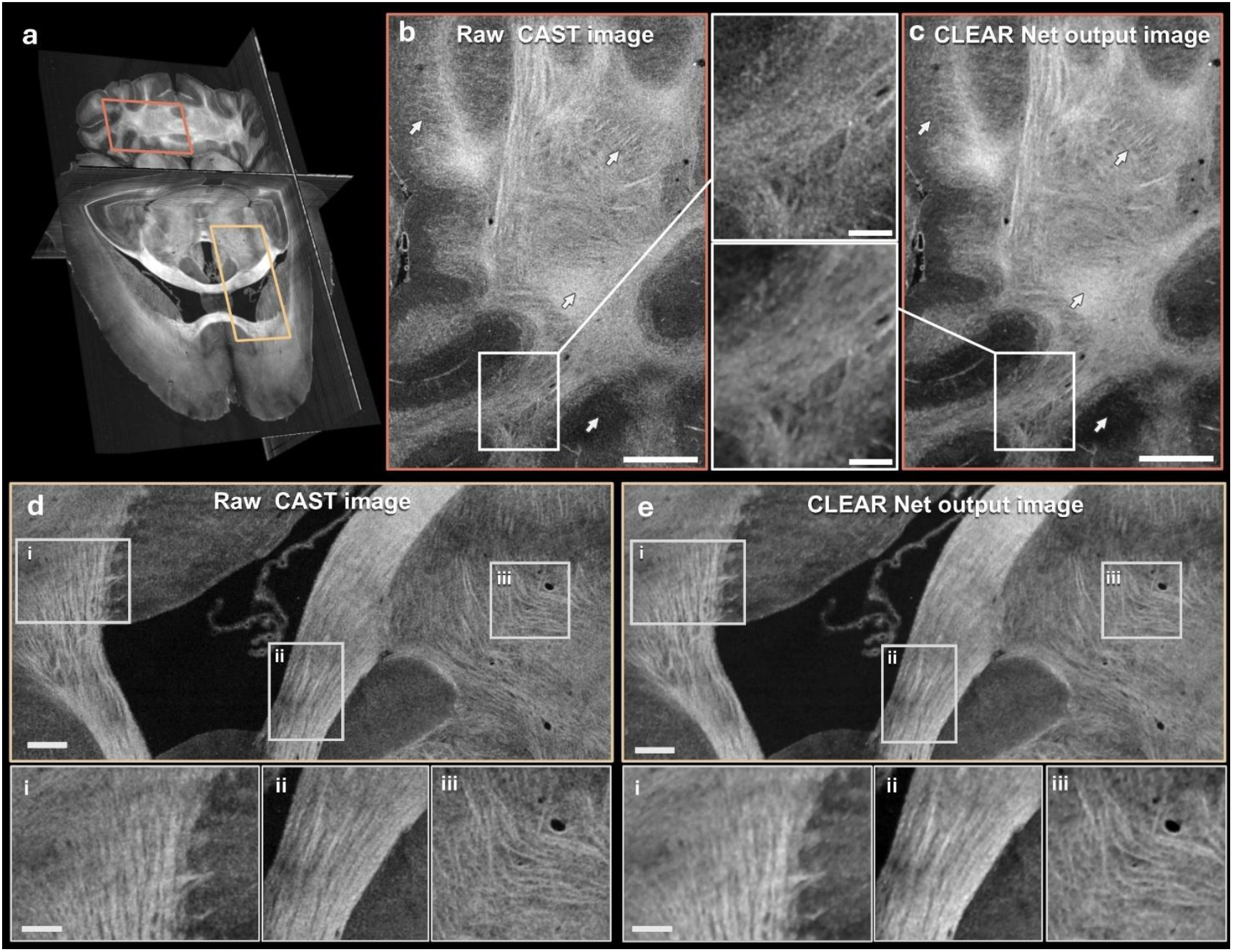
Representative qualitative results for the intact brain CAST dataset. (a) 3D rendering of the whole brain with two highlighted ROIs in the cerebellum (orange) and cerebrum (yellow). (b, c) Comparsion of raw CAST image with strong speckle noise and CLEAR Net speckle reduction image in the cerebellum. Scale bar, 500 μm. The white boxed regions in (b, c) show magnified views of fine structures with enhanced clarity in the generated images. Scale bar, 120 μm. (d, e) Comparsion of raw CAST image with strong speckle noise and CLEAR Net speckle reduction image in the cerebrum. Scale bar, 500 μm. Sub-ROIs scale bar, 120 μm.

### 3.2 Quantitative Evaluation and Comparison

We quantitatively compared CLEAR Net with other speckle reduction algorithms on CAST images. Figure 3 presents a comparison of speckle reduction results obtained using CLEAR Net, BM3D, PNLM, and Sub2Full on the corpus callosum and caudoputamen regions, together with an analysis of their corresponding frequency-domain characteristics. Fourier transforms were computed for green-box ROI regions to obtain the frequency-domain representations of each method. In the resulting FFT maps, the dashed circle denotes the cutoff frequency of CAST system (i.e., the wavelength at the full width at half maximum (FWHM)), while the solid white circle indicates the cutoff frequency of the images processed by each method. As shown in Fig.3b, the ground-truth images generated using the improved TNode algorithm exhibt effective speckle suprression compared with the raw CAST images, while preserving frequency characteristics consistent with the CAST system. In contrast, the higher-frequency components observed in the raw images primarily arise from speckle noise. the CLEAR Net outputs exhibit frequency characteristics closely matching those of the ground-truth images, with a cutoff frequency slightly higher than the ground truth. This indicates effective speckle reduction while maximally preserving true structural details. In contrast, BM3D yeild a large cutoff frequency than the ground truth, suggesting insufficient noise reduction of high-frequency nosie. PNLM produces a smaller cutoff frequency, reflecting excessive smoothing and loss of structural details. Sub2Full images display irregular FFT patterns, indicative of reconstruction artifacts and structural inconsistencies. Overall, the proposed approach achieves a better balance between speckle reduction and detail preservation, producing despeckled images with high visual similarity to the ground truth and superior frequency domain performance.

**Figure 3.**
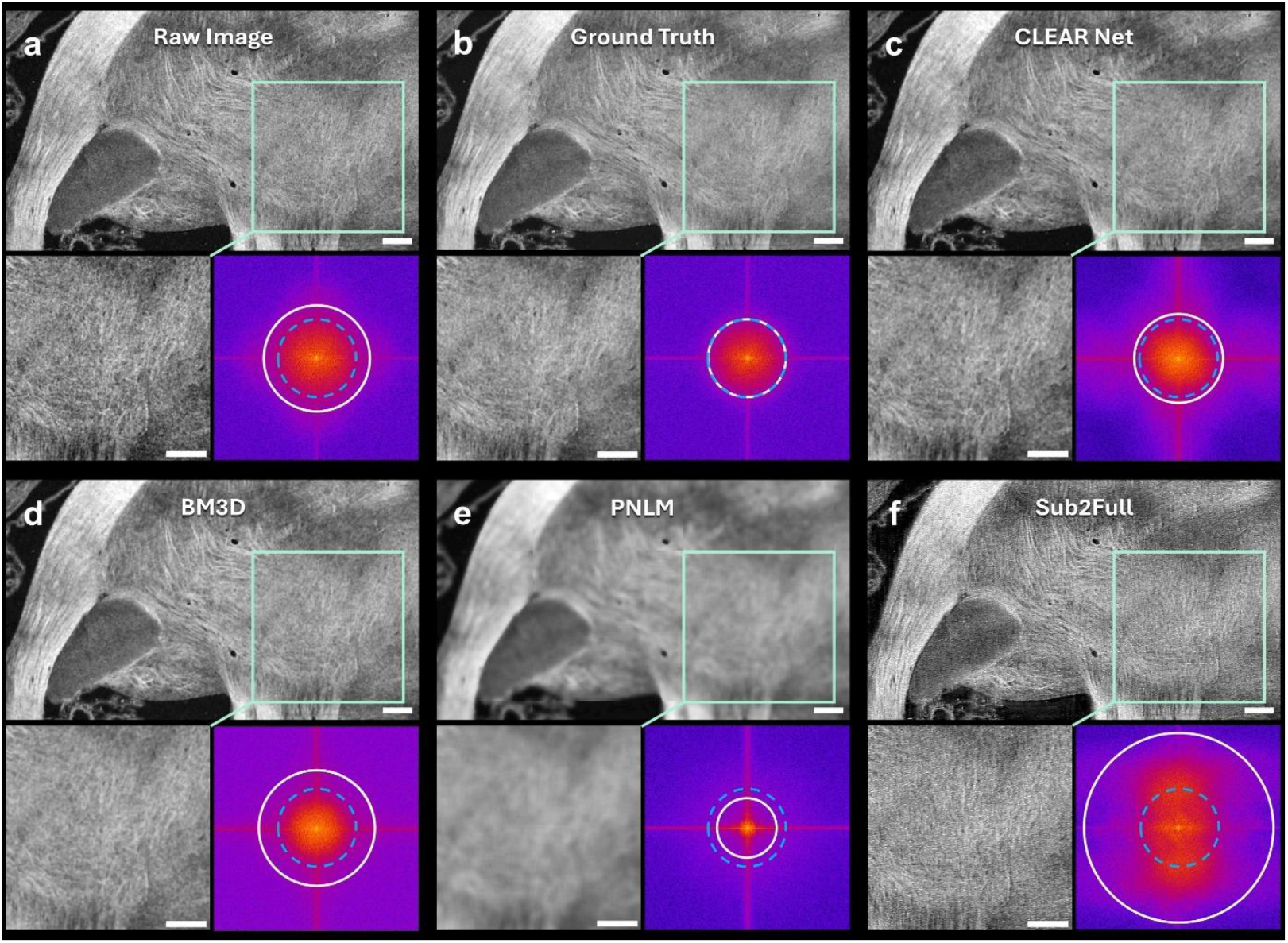
Comparison of speckle reduction performance on CAST images between CLEAR Net and other scattering-image despeckling algorithms. (a-f) Visualization of CAST images and magnified views of the corpus callosum and caudoputamen regions processed by different algorithms. (a) raw CAST image, (b) ground truth by improved TNode, (c) CLEAR Net output results, (d) BM3D result, (e) PNLM result, (f) Sub2Full result. The Fourier spectra of the magnified regions illustrate the ability of different methods to suppress noise while preserving fine structural details. The blue dashed lines indicate the cutoff frequency range of the CAST system, and the white circles indicate the corresponding image cutoff frequencies in the Fourier domain. Scale bar, 200 µm; ROI images, 250 µm.

**Figure 4.**
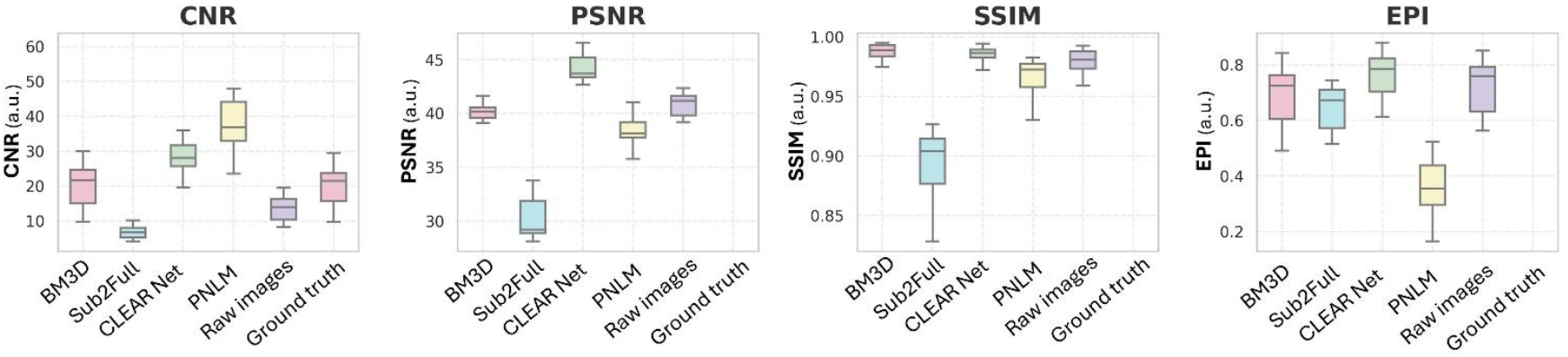
Quantitative comparison of speckle noise reduction performance among BM3D, PNLM, Sub2Full, and our CLEAR Net using four evaluation metrics: CNR, PSNR, SSIM, and EPI. PSNR, SSIM, and EPI were calculated within selected ROIs, while CNR was evaluated over the full image. The proposed method achieved the strong performance across all metrics, demonstrating superior noise reduction and structural preservation compared to existing approaches.

Furthermore, the quantitative comparison was performed between our proposed CLEAR Net and BM3D, PNLM, Sub2Full, as well as the nosisy input raw images, using four evaluation metrics: CNR, PSNR, SSIM, and EPI. PSNR, SSIM, and EPI were computed on randomly selected ROIs with the ground truth tomogram used as the reference image. CNR was evaluated across the entire image. For each image, six ROIs (each 512 × 512 pixels) were randomly chosen from twelve test slices for PSNR, SSIM, and EPI calculation. CNR values were obtained using a manually defined background region and multiple ROIs (each 128 × 128 pixels) through a custom-built graphical interface.

Across all four metrics, our method consistently outperformed the other despeckling approaches. In particular, the CLEAR Net results achieved the highest PSNR and EPI values, indicating superior noise reduction, improved edge preservation, and enhanced structural continuity. The SSIM of Gen was slightly lower than that of BM3D but remained comparable, while BM3D exhibited substantially lower PSNR and CNR, suggesting insufficient noise reduction despite its higher SSIM. Although PNLM achieved higher CNR, it showed extremely low EPI, reflecting excessive smoothing and loss of fine structural and edge details. In contrast, our method maintains a favorable balance between contrast enhancement and structural preservation. Collectively, these results demonstrate that our pipeline achieves superior performance in speckle reduction, structural integrity preservation, and overall image quality enhancement, providing a more consistent and reliable solution than traditional filtering methods and alternative deep learning-based approaches for high-fidelity tissue representation.

### 3.3 Fidelity Evaluation

Averaging multiple images acquired using hardware-based approaches with different speckle realizations can yield near speckle-free reference images; however, this requires a substantial increase in data volume, typically exceeding 20 acquisitions to achieve significant speckle reduction. For whole-brain CAST imaging, such averaging would result in datasets larger than 160 TB, rendering both data acquisition and reconstruction times impractical. Therefore, the improved TNode-processed images were used as ground-truth images for training CLEAR Net. To validate the fidelity of CLEAR Net outputs, we introduced a diffuser into the system to generate independent speckle realizations and used the averaged results as a gold-standard reference for comparison with the network outputs (Fig.5a).

**Figure 5.**
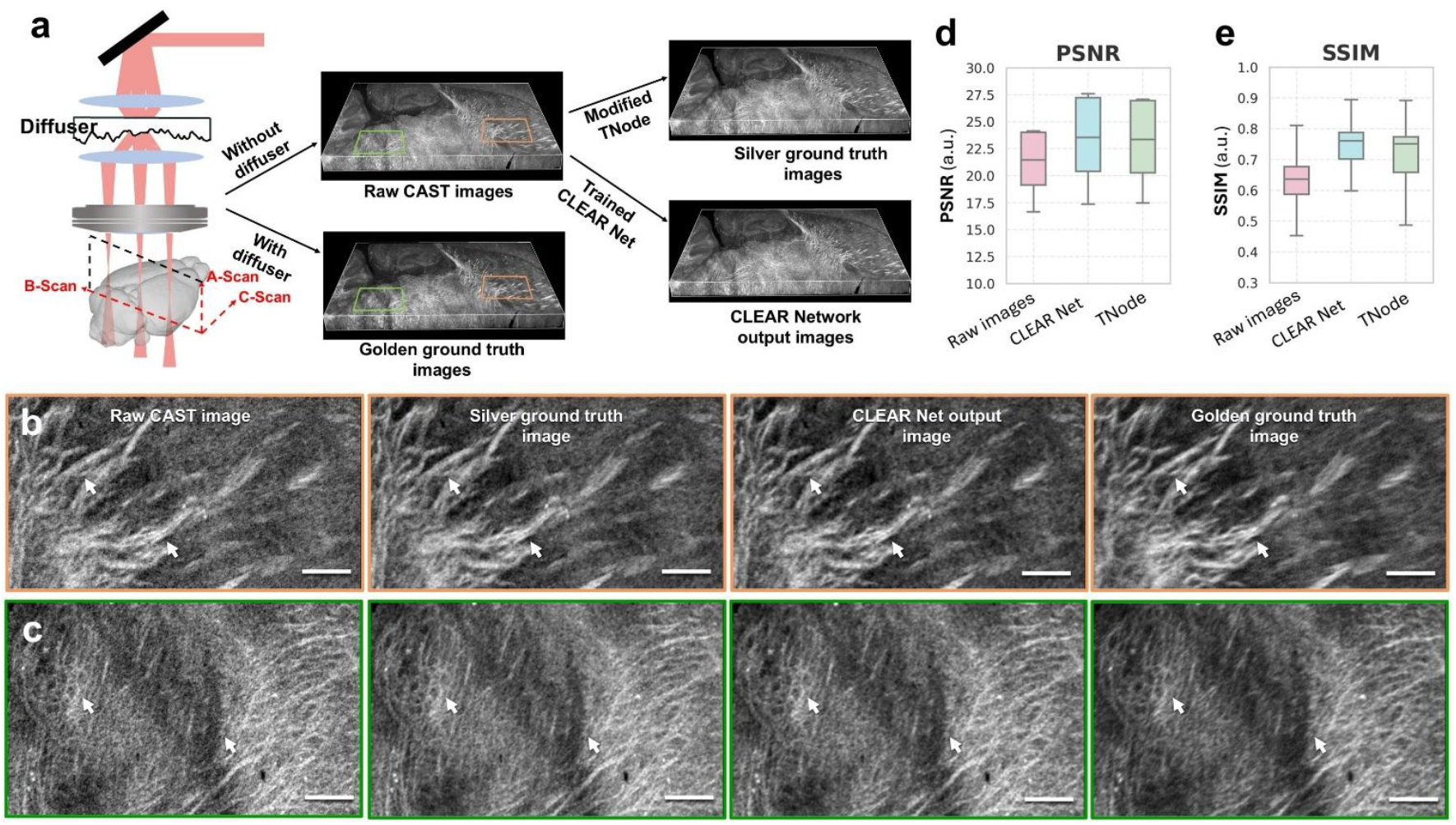
Quantitative comparison between CLEAR Net and the hardware-based speckle modulation-derived golden ground truth validates the fidelity of the proposed method. (a) Illustration of image acquisition and processing pathways. (b-c) Qualitative comparison in a fine-detail region containing white matter structures. Scale bar, 100 µm. (d, e) PSNR and SSIM comparison of raw input, TNode, and network output computed on 200 randomly selected 512 × 512 patches through half-brain dataset, using the golden ground truth as the reference.

The TNode-based results were generated using the same algorithmic parameters as those used to produce the training ground truth. Network outputs were obtained by directly applying the trained model to new mouse brain data without fine tuning. The golden ground truth was generated by averaging 22 repeated acquisitions at the same sample position, speckle noise was effectively suppressed while preserving underlying structures. Owing to the time-intensive nature of this procedure, only a subset of the brain volume was acquired. In this validation, the Golden GT served as an independent reference, whereas network training relied exclusively on algorithmically generated ground truth.

The speckle reduction results of CLEAR Net in gray and white matter regions were separately compared with the gold-standard reference (Fig. 5b-c). As shown in Fig. 5b, the input image is dominated by speckle noise that obscures underlying tissue signals, whereas the gold-standard reference (Golden GT) exhibits substantial speckle reduction achieved through diffuser-based acquisition. Both TNode and CLEAR Net effectively suppress speckle and improve image quality. Notably, CLEAR Net provides more uniform noise reduction while better preserving image texture, resulting in closer visual agreement with the Golden GT. Focusing on a fine-detail white matter region (Fig. 6c), speckle noise in the input image disrupts structural continuity and leads to fragmented textures. While both TNode and CLEAR Net recover overall white matter continuity, the network output reveals more coherent and continuous fibrous patterns, with finer structural delineation that more closely resembles the Golden GT.

**Figure 6.**
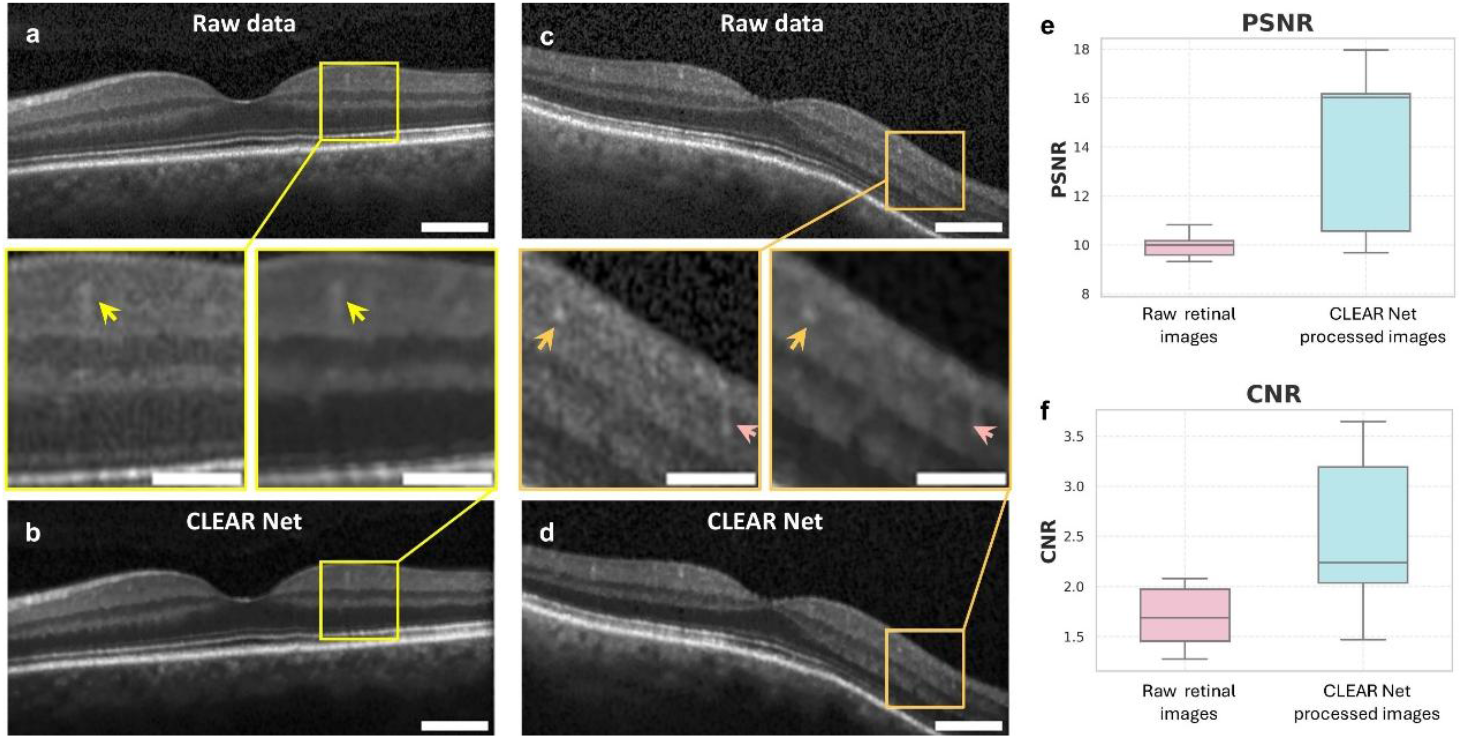
Performance of CLEAR Net on the publicly available retinal dataset. (a, c) Raw retinal OCT images. (b, d) Corresponding outputs generated by CLEAR-S3 net. Yellow and orange boxes indicate zoomed-in regions, and arrows highlight preserved structural details. Scale bar, 70 µm; ROI images, 30 µm. (e) PSNR comparison between raw images and generated images. (f) CNR comparison between raw images and generated images.

Quantitative evaluations using the gold-standard reference are shown in Fig. 6d–e. Both TNode and CLEAR Net achieve higher PSNR and SSIM than the raw input, with CLEAR Net consistently outperforming TNode. The close agreement between TNode results and the hardware-based reference supports the use of TNode-generated images as surrogate ground truth for large-scale training. Moreover, the strong correspondence between the network output and the gold-standard reference indicates that CLEAR Net effectively learns and preserves the key structural characteristics of the underlying signal.

### 3.4 Generalizability Evaluation

To assess the generalization capability of the proposed pipeline beyond brain images, we directly applied the trained network to the Kermany Retinal OCT Dataset^26^ without any fine-tuning. Figure 6 compares raw retinal OCT images with their corresponding outputs. The zoomed-in regions (yellow and orange boxes) show that speckle noise is effectively suppressed, producing visibly cleaner backgrounds and clearer depiction of retinal layers. Generated images also display enhanced contrast, which sharpens layer boundaries and improves overall interpretability. In addition, the continuity of the layer structures is strengthened, with transitions appearing smoother and less affected by noise. These improvements are achieved while maintaining fine anatomical details within the retina. The performance gains are further confirmed by PSNR and CNR measurements (Fig. 6e-f). The generated images exhibit a significant increase in PSNR and CNR, indicating improved fidelity and clearer contrast separation under reduced noise levels.

Both qualitative and quantitative evidence consistently show that the method generalizes well to retinal imaging domain. The consistent qualitative and quantitative improvements on retinal OCT images achieved without any additional fine-tuning demonstrate that our pipeline maintains its performance across anatomical domains. These findings indicate that the method possesses meaningful generalization capability, supporting its applicability to a broader range of coherent imaging scenarios.

## 4. Conclusion

This study proposes a machine learning-based speckle reduction pipeline for CAST brain imaging. By leveraging high-fidelity references, volumes generated with an improved TNode algorithm, and through the design of network architecture and loss functions, the proposed approach achieves a balance between speckle reduction and fine structural preservation, establishing an effective processing pipeline tailored for CAST brain imaging.

Experiments on CAST mouse brain datasets demonstrate that the method can generate despeckled images close to the ground truth, clearly preserving layered structures and fine textures while maintaining overall volume consistency. Compared with the original images, the generated results are visually clearer, speckle noise is effectively suppressed, and quantitative metrics also show significant improvements in overall image quality. Existing deep learning-based despeckling pipelines fail to generalize well to CAST brain data, whereas our proposed framework produces results that are both visually and quantitatively superior. Compared with conventional approaches such as BM3D and PNLM, our method not only achieves stronger noise reduction but also better preserves fine anatomical details, including white matter structures. While ensuring high fidelity and clarity, the computational efficiency is significantly higher than that of traditional iterative methods, providing a feasible pathway for large-scale CAST brain imaging data processing. Moreover, the framework demonstrates a certain degree of generalizability across different tissue structures and organ types, maintaining high-quality despeckling beyond the original training domain.

The proposed method generates low-noise, structurally faithful images that provide more reliable inputs for downstream tasks such as automated segmentation and feature extraction, enabling accurate capture of details that are otherwise prone to speckle interference and thereby improving the precision of subsequent analyses. In neuroscience research and pharmacological intervention studies, this approach also facilitates more accurate quantification of subtle structural changes while reducing the impact of speckle on experimental outcomes.

In future work, we will extend this method to other tissues and imaging systems and develop more robust, multi-task models. In addition, we plan to explore models that explicitly learn the statistical properties of speckle independent of structural content, enabling a unified despeckling framework applicable across coherent imaging modalities, thereby further advancing the application of coherent imaging in broader biomedical research.

## Acknowledgements

This research was funded by National Institutes of Health (NIH) (R00AG059946)

